# Combined degradome and replicated small RNA sequencing identifies *Brassica napus* small RNAs responsive to infection by a necrotrophic pathogen

**DOI:** 10.1101/2020.11.20.392209

**Authors:** Roshan Regmi, Toby E. Newman, Lars G. Kamphuis, Mark C. Derbyshire

## Abstract

**Background:** Small RNAs are short non-coding RNAs that are key gene regulators controlling various biological processes in eukaryotes. Plants may regulate discrete sets of sRNAs in response to pathogen attack. *Sclerotinia sclerotiorum* is an economically important pathogen affecting hundreds of plant species, including the economically important oilseed *Brassica napus*. However, there are limited studies on how regulation of sRNAs occurs in the *S. sclerotiorum* and *B. napus* pathosystem.

**Results:** We identified different classes of sRNAs from *B. napus* by high throughput sequencing of replicated mock and infected samples at 24 hours post-inoculation (HPI). Overall, 3,999 sRNA loci were highly expressed, of which 730 were significantly upregulated during infection. Degradome sequencing identified numerous likely sRNA targets that were enriched for immunity-related GO terms, including those related to jasmonic acid signalling, during infection. A total of 73 conserved miRNA families were identified in our dataset. Degradome sequencing identified 434 unique cleaved mRNA products from these miRNAs, of which 50 were unique to the infected library. A novel miR1885-triggered disease resistance gene-derived secondary sRNA locus was identified and verified with degradome sequencing. We also experimentally validated silencing of a plant immunity related ethylene response factor gene by a novel sRNA using 5’-RACE.

**Conclusions:** The findings in this study expand the framework for understanding the molecular mechanisms of the *S. sclerotiorum* and *B. napus* pathosystem at the sRNA level.

## Background

Small RNAs (sRNAs) are short non-coding RNAs, ranging in size from 18-30 nt, that are important for gene expression regulation and genome stability in eukaryotes [1]. There are three major sRNA classes, microRNAs (miRNAs), short interfering RNAs (siRNAs) and P-element Induced WImpy (PIWI) associated RNAs (piRNAs); while the latter only occur in animals [2], the former two are found in plants.

Different types of sRNAs have different biogenesis pathways [3]. The canonical miRNA pathway comprises several steps. Long precursor genes are transcribed by RNA polymerase II (Pol II). These several hundred base pair long sequences fold back to form hairpin-like structures commonly called pri-miRNAs. RNAses called Dicer like (DCL) proteins precisely cleave the pri-miRNAs to form duplexes.

Plants have many classes of siRNA [1, 4–6]. The main siRNA classes are the hairpin-siRNAs (hp-siRNAs), natural-antisense siRNAs (natsiRNAs), secondary siRNAs and heterochromatic siRNAs (hetsiRNAs) [1]. These classes have distinct biogenesis pathways and may involve DCL protein mediated cleavage of duplexes from either very long hairpins (hp-siRNAs) or double stranded RNAs generated from single stranded precursors by RNA dependent RNA polymerase (RDR) enzymes [1].

Small RNAs in plants target genes for silencing through the RNA interference pathway. One of the strands of the sRNA duplex produced by DCL cleavage of the precursor has a lower thermodynamic stability at the 5’ end and is loaded into a protein in the Argonaute (AGO) family. This strand, known as the guide strand, targets sequences for silencing through complementary base pairing. The other strand, known as the passenger strand (sRNA*), is often quickly degraded but may also be involved in silencing of complementary sequences. AGOs form a complex with other proteins known as the RNA-induced silencing complex (RISC) to facilitate gene silencing [7]. Once the sRNA finds its complementary targets, the RISC suppresses their expression. The targets of miRNAs are normally complementary mRNA transcripts, and repression of gene expression may be mediated by inhibition of translation or mRNA cleavage. The siRNAs regulate gene expression by both RNA-directed methylation of complementary DNA and mRNA cleavage [1].

In addition to silencing mRNAs, miRNA-mediated cleavage of mRNAs or non-coding RNA precursors may also produce secondary siRNAs. These are often described as phased siRNAs (pha-siRNAs) as they appear at precise 21-22 nucleotide intervals from the miRNA cleavage site. Loci that produce pha-siRNAs are known as ‘PHAS’ loci [8]. The secondary RNAs produced by PHAS loci may silence the gene from which they are derived or they may act in *trans* to silence the expression of other genes; the latter type of secondary siRNAs are known as trans-acting siRNAs (ta-siRNAs) and their biogenesis loci are often referred to as ‘TAS’ genes.

While a large number of miRNA-triggered secondary siRNAs have been identified in the genomes of plants, only a few have been experimentally validated [9–11][12–14]. Four miRNA triggered ta-siRNA families have been characterized in the model plant *Arabidopsis thaliana* [11]. Of these, the miRNA390-triggered TAS3 genes were found to be conserved across various plant species.

The formation of pha-siRNAs depends on several protein components, including SUPPRESSOR OF GENE SILENCING 3 (SGS3), RDR6 and DCL4 [11]. Studies on PHAS loci in different plants have shown that miRNAs trigger pha-siRNA production from many types of transcript, including noncoding RNAs, and the mRNAs of disease resistance and pentratricopetide repeat genes [10]. Nucleotide-binding site leucine-rich repeat (NBS-LRR) genes form the largest set of genes identified so far that can potentially produce pha-siRNAs upon binding of specific miRNAs [15].

During viral infection of plants, changes in the accumulation of miRNAs result in production of different pha-siRNAs [16]. In legumes and tomato, a number of miRNA families are involved in triggering pha-siRNAs by binding to the transcripts of NB-LRR genes [10, 17]. In tomato, the abundance of secondary siRNAs from disease resistance genes was lower during bacterial and viral infection, suggesting that pha-siRNA production is important for fine-tuning defence responses [17]. Recently Cui et al. (2020) demonstrated the role of miR1885-mediated ta-siRNA expression in maintaining plant growth and immunity in *B. napus* upon viral infection [18].

The roles of pha-siRNAs in plant response to bacterial and viral infection have been investigated in several studies but little is known about their roles in responding to pathogenic fungi. One of the few studies on this subject was by Wu et al. (2017). This study characterized pha-siRNAs produced by tomato in response to *Botrytis cinerea* infection [12]. It was found that many pathogen-responsive tomato pha-siRNAs downregulate transcription factors, which is suggestive of a broad role in the regulation of gene expression regulation [14].

Other than pha-siRNAs, plants under pathogen attack may employ various sRNA-regulated immune pathways [19, 20]. For example, while studying the sRNAome in wheat cultivars during *Puccinia graminis* infection, Gupta et al. (2012) reported that miR408 exhibits different expression patterns in susceptible and resistant cultivars after a two-day course of infection [21].

Some immunity-related sRNAs have also been functionally characterised. For example, in the model plant *A. thaliana*, microRNA393 targets different auxin signalling genes to confer antibacterial resistance [22] and miR408 is a negative regulator of plantacyanins and laccase genes [23]. These latter genes have roles in stress responses, cell-to-cell signalling and maintaining plasticity and vigour of the cell wall. In addition, overexpression of miR7695 results in an incremental increase in resistance in rice against the blast fungus *Magnaporthe oryzae* [24].

Canola (*Brassica napus*) is an economically important oilseed crop grown worldwide [25]. Sclerotinia stem rot (SSR), caused by the fungus *Sclerotinia sclerotiorum*, is an important disease that causes large economic losses in canola [26]. Some studies have been conducted in *Brassica* spp. to identify plant-specific miRNAs [27–29] under biotic and abiotic stresses. There have been two studies on *B. napus* miRNA expression upon *S. sclerotiorum* infection [30, 31]. However, these studies were performed with a single sRNA library at 12 and 48 hours post-inoculation (HPI) without any replicates, so differential expression analysis was not possible.

In comparison to mature miRNAs deposited in miRBase for other plants like *Medicago truncatula*, *O. sativa* and *A. thaliana,* the number of miRNAs for *B. napus* is quite low, suggesting many miRNAs in *B. napus* are yet to be discovered. Furthermore, little is known about the triggers of PHAS loci in the *B. napus* genome and their functions in gene regulation. Therefore, to assess differential expression of sRNAs, identify new pathogen-responsive miRNAs and characterise the role of secondary sRNAs in the *B. napus* response to *S. sclerotiorum*, we developed replicated sRNA libraries for mock-inoculated and *S. sclerotiorum* inoculated leaves 24 hours post treatments to characterize different classes of sRNAs. To identify targets of these sRNAs, we also performed degradome sequencing, 5’-rapid amplification of cDNA ends (5’-RACE) and quantitative PCR (qPCR).

## Results

### Overview of sequencing results

To determine the role of *B. napus* sRNAs during *S. sclerotiorum* infection we sequenced six sRNA libraries on the Illumina platform from three replicates each of mock and infected samples at 24 HPI when SSR symptoms manifested on leaves. A total of 152,090,773 raw reads were obtained from the six libraries. We retained 126,887,984 (83.24 %) high quality reads after adapter trimming and length filtering (18-30 nt) from these six libraries (Table 1). Assignment and removal of ambiguous reads (that map to both plant and fungal genome) resulted in 41,797,278 unique *B. napus* sRNA reads that match best to the *B. napus* genome across all libraries. The reads that potentially originated from structural RNAs (rRNAs, snRNAs, snoRNAs) accounted for ~ 5 % of this total. The clean, high-quality mappable reads were then aligned to the *B. napus* genome. The overall alignment rate was 88.7 % with the highest percentage mapping in the mock samples (above 98 %), while in infected samples an average of ~ 78.9 % of reads mapped, ranging from 77 to 86.3 % between replicates. Among the mapped reads, ~ 86 % were mapped to more than one genomic locus revealing that these sRNAs originated from genomic repeats. From our dataset, we found 14 % of sRNA reads that uniquely mapped to a single genomic locus. Table 1 provides an overall summary of the sequencing data.

**Table 1.**
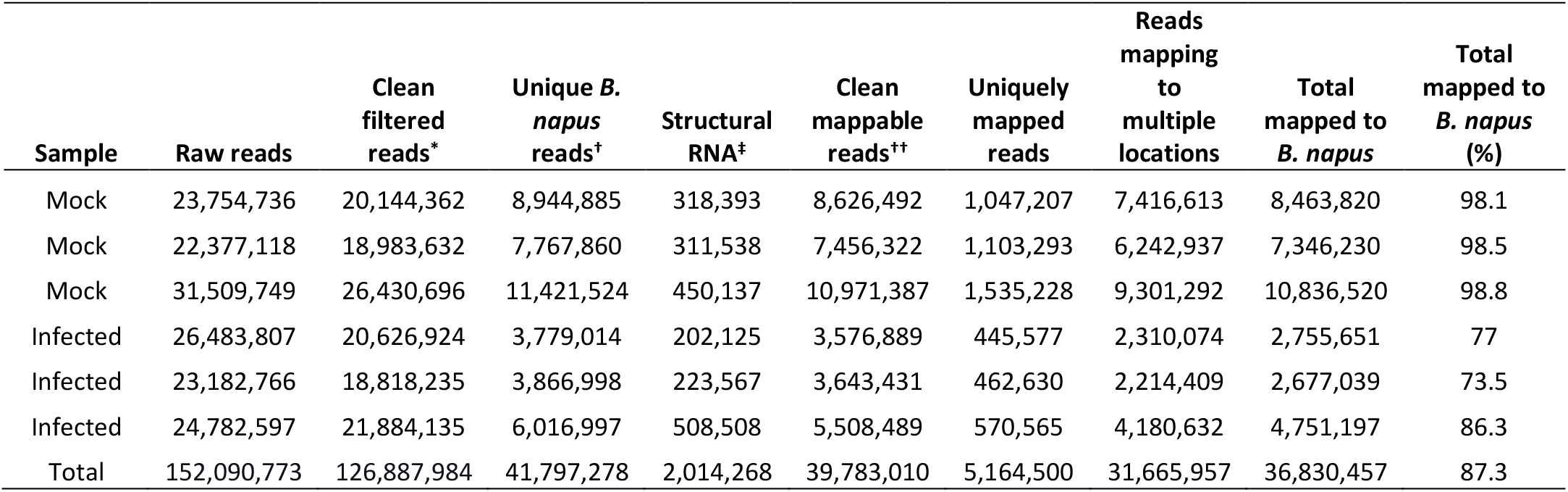
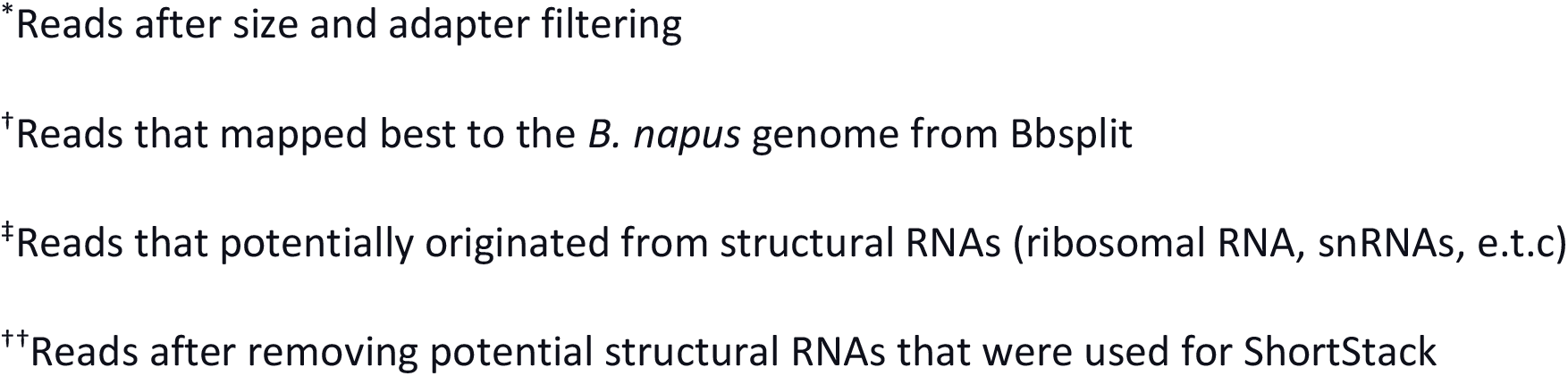
An overall summary of the sequencing data.

| Sample | Raw reads | Clean filtered reads* | Unique *B. napus* reads<sup>†</sup> | Structural RNA<sup>‡</sup> | Clean mappable reads<sup>††</sup> | Uniquely mapped reads | Reads mapping to multiple locations | Total mapped to *B. napus* | Total mapped to *B. napus* (%) |
| --- | --- | --- | --- | --- | --- | --- | --- | --- | --- |
| Mock | 23,754,736 | 20,144,362 | 8,944,885 | 318,393 | 8,626,492 | 1,047,207 | 7,416,613 | 8,463,820 | 98.1 |
| Mock | 22,377,118 | 18,983,632 | 7,767,860 | 311,538 | 7,456,322 | 1,103,293 | 6,242,937 | 7,346,230 | 98.5 |
| Mock | 31,509,749 | 26,430,696 | 11,421,524 | 450,137 | 10,971,387 | 1,535,228 | 9,301,292 | 10,836,520 | 98.8 |
| Infected | 26,483,807 | 20,626,924 | 3,779,014 | 202,125 | 3,576,889 | 445,577 | 2,310,074 | 2,755,651 | 77 |
| Infected | 23,182,766 | 18,818,235 | 3,866,998 | 223,567 | 3,643,431 | 462,630 | 2,214,409 | 2,677,039 | 73.5 |
| Infected | 24,782,597 | 21,884,135 | 6,016,997 | 508,508 | 5,508,489 | 570,565 | 4,180,632 | 4,751,197 | 86.3 |
| Total | 152,090,773 | 126,887,984 | 41,797,278 | 2,014,268 | 39,783,010 | 5,164,500 | 31,665,957 | 36,830,457 | 87.3 |

\*Reads after size and adapter filtering
<sup>†</sup>Reads that mapped best to the *B. napus* genome from Bbsplit
<sup>‡</sup>Reads that potentially originated from structural RNAs (ribosomal RNA, snRNAs, e.t.c)
<sup>††</sup>Reads after removing potential structural RNAs that were used for ShortStack

To determine the grouping of infected and mock libraries we performed a principal component analysis on mean normalized counts from DESeq2 (Fig 1A). The principal component analysis showed the replicated datasets were well grouped for two treatment groups, i.e. mock and infected, suggesting large overall differences between these treatments. Mock and infected samples were separated along principal component 1, which explained 99 % of the variance. There was some spread between the infected samples along principal component 2. However, variance between these samples along this axis is negligible, since only 1 % of the variance was explained by PC2.

**Figure 1.**
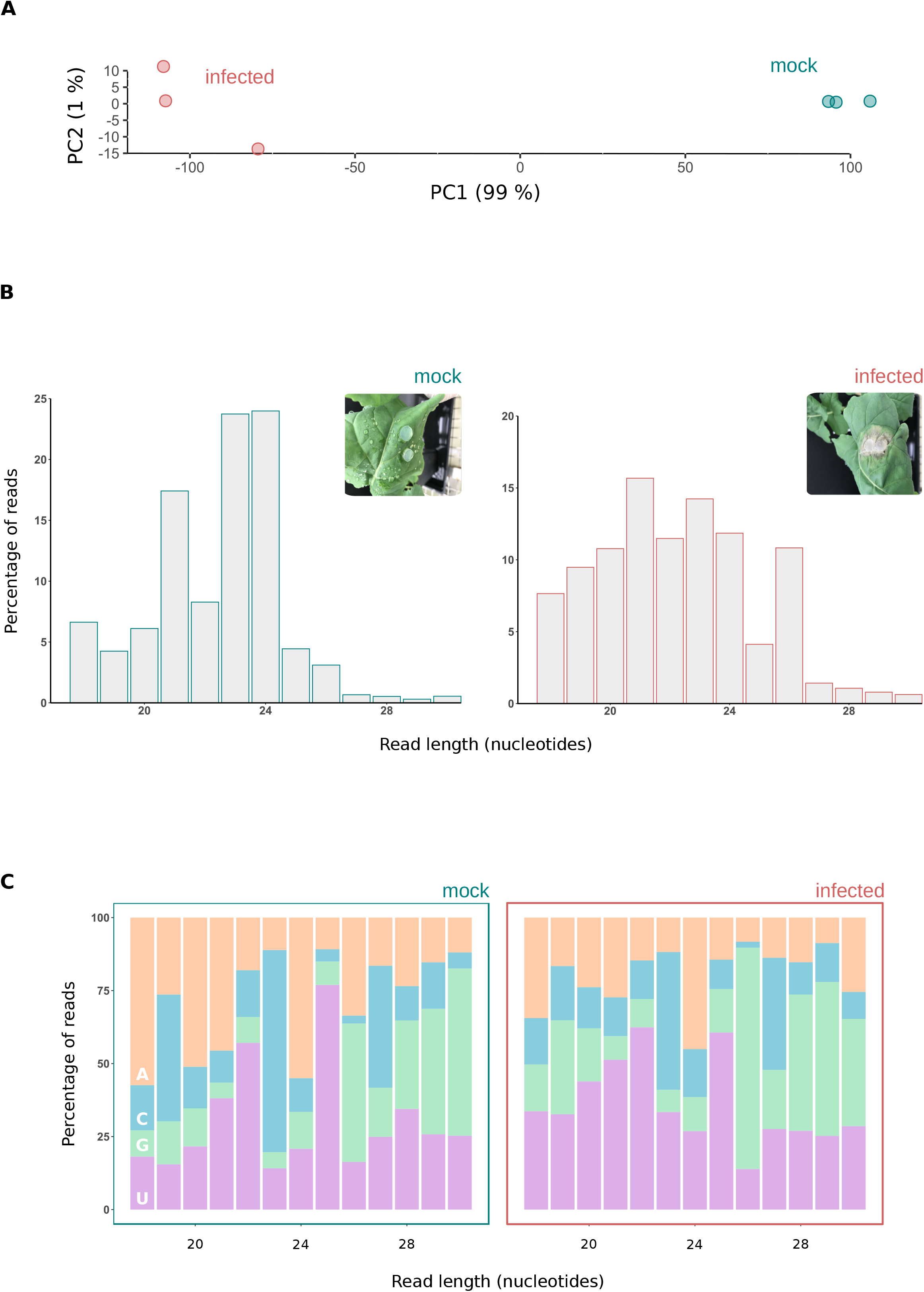
Changes in size class and 5’ nucleotide of *Brassica napus* sRNAs in response to *Sclerotinia sclerotiorum* infection. **(A)** A principal component analysis based on normalized read counts from DESeq2. The x axis shows principal component 1, which explained 99 % of the variance, and the y axis shows principal component 2, which explained 1 % of the variance. The infected samples are depicted with red circles and the mock samples depicted in turquoise. **(B)** Histogram of read sizes from pooled mock and infected samples. Inset: a representative picture of *Brassica napus* leaves under both treatments (24 hours post-inoculation (HPI)). Left: for the pooled replicates of the mock sample, y axis depicts the percentage of reads across all three replicates and the x axis read length in nucleotides. Right: for the pooled replicates of the infected sample, same information as for the mock sample. **(C)** Results from pooled replicates for the mock (left) and infected (right). Showing percentage of reads (y axis) of each size class (x axis) that had each of the four nucleotides (AGCU) in their 5’ position.

### Characteristic features of the *Brassica napus* small RNA population

To determine which sRNAs were induced in response to infection, we produced three sequencing replicates each from mock and infected samples. The following metrics are based on the pooled biological replicates for each treatment, mock and infected. Size class distribution and 5’ nucleotide bias are two important characteristics to determine the origin and activities of sRNAs. To determine whether there may be a difference in the composition of sRNA origins in mock and infected samples, we analysed the nucleotide length and 5’ nucleotide bias of these sRNAs. Interestingly, we found a difference in length distribution between mock and infected samples (Fig 1B), suggesting that upon infection, sRNA biogenesis mechanisms are altered. In mock samples, almost 50 % of total reads belonged to 24 and 23 nucleotide (nt) sRNAs followed by 21 nt. Adenine was enriched as the 5’ nucleotide in 24 nt sRNAs while cytosine was more abundant in 23 nt sRNAs. A 5’ nucleotide bias toward uracil was present mostly in 22 nt sRNAs (Fig 1C).

In infected samples, size classes were more uniform than in mock samples (Fig 1B). The most abundant read size was 21 nt with a slight 5’ uracil bias, followed by 23 nt with a slight cytosine bias (Fig 1C). We also found a peak in 26 nt in infected samples with a 5’ guanine bias. Similarly, size classes of 18, 19, 20 and 22 nt were also more abundant in infected samples. Previous reports presented similar data with a 5’ uracil bias in 21 nt and a 5’ adenine bias in 24 nt sRNAs. The 24 nt 5’ adenine biased siRNAs have been previously shown to be involved in RNA dependent DNA methylation in *A. thaliana* with preferential loading into AGO4, while the 21 nt 5’ uracil biased sRNAs have preferential loading into AGO1 [51]. Overall, our results suggest a marked shift in the types of sRNAs expressed from mock to infected *B. napus* leaves.

### A total of 730 unique *Brassica napus* small RNAs are upregulated in response to *Sclerotinia sclerotiorum* infection

To assess what *B. napus* sRNAs accumulate in response to infection with *S. sclerotiorum*, we performed a differential expression analysis with DESeq2. We did not only consider differential expression of miRNAs but the entire sRNA-ome in *B. napus*. ShortStack predicted 121,977 sRNA loci, 104,421 of which were likely Dicer-derived. Among these loci, 3,999 were highly expressed, with at least 100 raw major RNA sequencing reads (Supplementary Table 2). If these sRNAs were responding to infection, we hypothesised that they might be more expressed in infected samples as compared to mock samples. We found 915 loci significantly altered in their expression in infected samples compared to the mock samples. Among these, 730 were upregulated in *B*. *napus* after *S. sclerotiorum* infection; these loci produced 565 unique sRNAs based on the major sRNAs predicted by ShortStack (Fig 2A).

**Figure 2.**
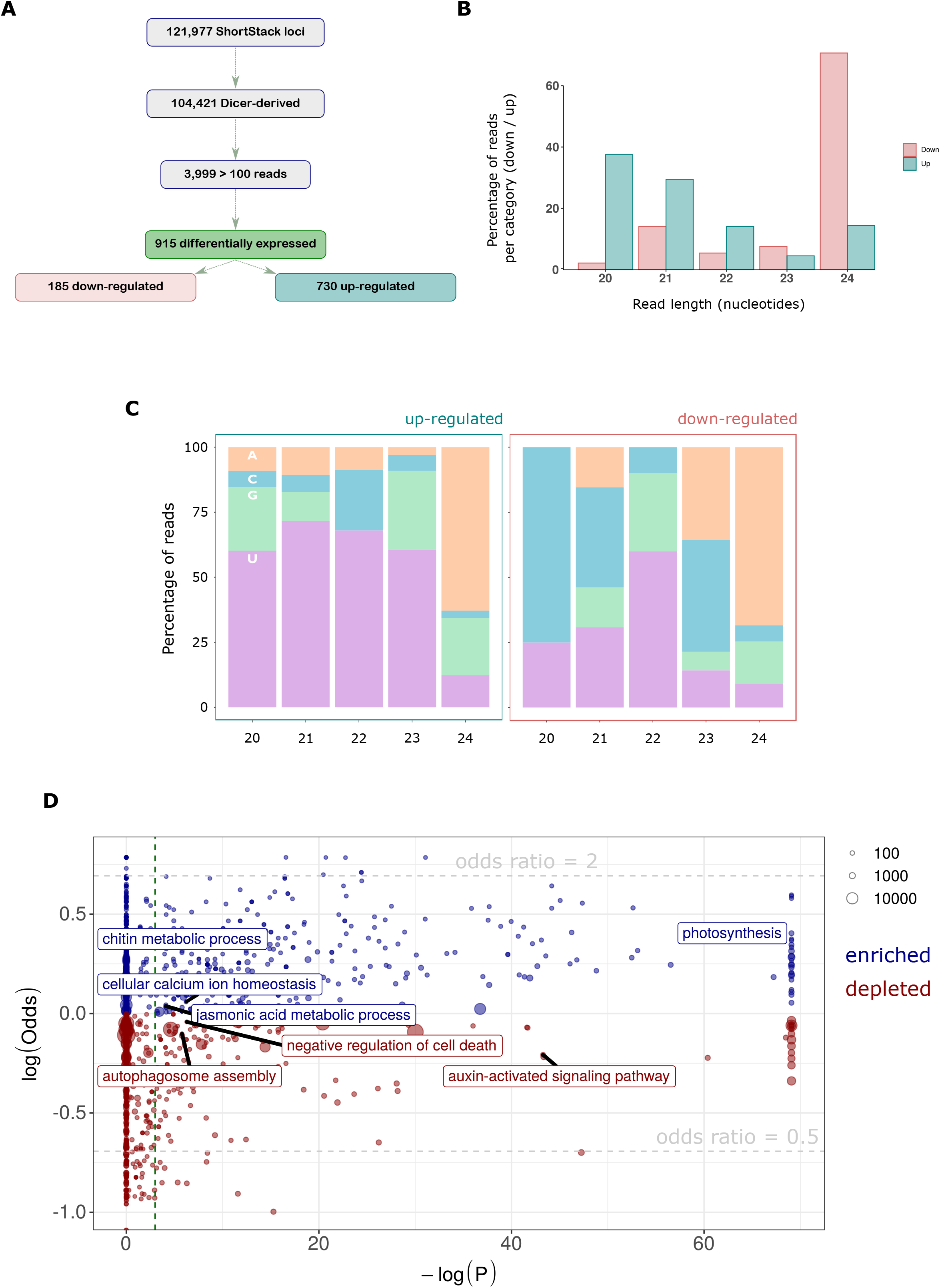
Small RNA population identified from *Brassica napus* genome in response to *Sclerotinia sclerotiorum* infection. **(A)** A flow chart showing total, Dicer-derived and highly expressed loci with a major RNA reads of => 100 reads, as predicted by ShortStack, and differentially expressed loci identified from DESeq2. **(B)** Histogram of read sizes from upregulated and downregulated sRNA loci, y axis depicts the percentage of reads in each category (upregulated or down-regulated) and the x axis read length in nucleotides. **(C)** Histogram of read sizes showing percentage of reads (y axis) of each size class (x axis) that had each of the four nucleotides (AGCU) in their 5’ position for upregulated loci (left) and downregulated loci (right). **(D)** Degradome targets of *Brassica napus* sRNAs in response to *Sclerotinia sclerotium* infection. Significance of the enrichment test (–log(*p*)) is on the x axis; the dashed vertical line is a *p* value of 0.05. The logarithm of the ratio of the observed to expected proportions of the GO terms in sRNA targets is on the y-axis. The size of points represents the number of degradome sRNA target candidates annotated with the GO term. The blue points are GO terms enriched in the infected sample and those in red are depleted in the infected sample. Some terms discussed in the text are highlighted. These terms showed enrichment in the infected sample and depletion in the mock (blue) or vice versa (red).

These 730 upregulated sRNAs were mostly enriched for 20 and 21 nt sequences, while the 185 downregulated major sRNAs were enriched for 24 nt sequences (Fig 2B). Uracil was enriched at the 5’ ends of all the size classes for upregulated sRNAs except 24 nt, which had a 5’ adenine bias (Fig 2C). However, downregulated sRNAs exhibited a 5’ cytosine bias at 20 nt and 21 nt (Fig 2C). Size classes 22 and 24 nt shared a common 5’ bias of uracil and adenine respectively in both sample groups. Overall, our data add weight to the hypothesis that sRNA classes with distinct biogenesis and targeting pathways were expressed in response to *S. sclerotiorum* challenge.

### Jasmonic acid and auxin signalling are regulated by small RNAs in *Brassica napus* during *Sclerotinia sclerotiorum* infection

To gain a global overview of the *B. napus* sRNA regulatory cascade we used all sequenced non-redundant sRNA reads to find their targets using degradome tags in mock and infected libraries. From 5,169,927 valid non-redundant plant sRNA reads, 311,588 and 183,056 total cleavage products with 301,246 and 44,359 unique tags were found from the mock and infected samples, respectively. The redundancy in targets suggested that one gene may be regulated by multiple sRNAs [52]. We identified multiple Biological Process GO terms both enriched (over-abundant) and depleted among sRNA targeted genes in the infected sample (Supplementary Table 3). These included terms such as ‘glycolytic process’ (GO:0006096), ‘mRNA splicing, via spliceosome’ (GO:0000398), ‘protein kinase C-activating G protein’ (GO:0007205), ‘leucine biosynthetic process’ (GO:0009098), ‘methylation’ (GO:0032259), ‘DNA replication’ (GO:0006260) and ‘microtubule-based movement’ (GO:0007018).

Several terms were significantly enriched in mock samples and significantly depleted in infected samples, and vice versa (Fig 2D). The terms ‘chitin metabolic process’ (GO:0006032), ‘cellular calcium ion homeostasis’ (GO:0006874), ‘jasmonic acid metabolic process’ (GO:0009694) and ‘photosynthesis’ (GO:0015979) were enriched in infected samples and depleted in mock samples. The terms ‘negative regulation of cell death’ (GO:0060548), ‘autophagosome assembly’ (GO:0000045) and ‘auxin-activated signalling pathway’ (GO:0009734) were depleted in the infected samples and enriched in mock samples. These data suggest that, among other things, jasmonic acid signalling is repressed by plant sRNAs after 24 hours of infection, whereas auxin signalling is de-repressed.

We also used the degradome sequencing data to investigate targets of sRNAs upregulated during infection. A total of 64 target genes were identified from upregulated sRNAs this way (Table 2). The representative T-plots for four of these genes that were identified in infected libraries, which were assigned to different PARESnip2 confidence categories are presented in Fig 3. Among these 64 targets, 10 were found in both libraries resulting in 29 and 15 unique targets for mock and infected samples, respectively. These negatively regulated genes induced upon infection were annotated with transcription factor-related InterPro terms such as ‘ABC transcription factor’, ‘heat shock response’ and disease resistance protein-like ‘Zinc finger l domain’ and ‘leucine zipper domain’. This suggests that *B. napus* transcriptional regulatory networks may be modified by sRNAs specifically induced during infection.

**Table 2.**
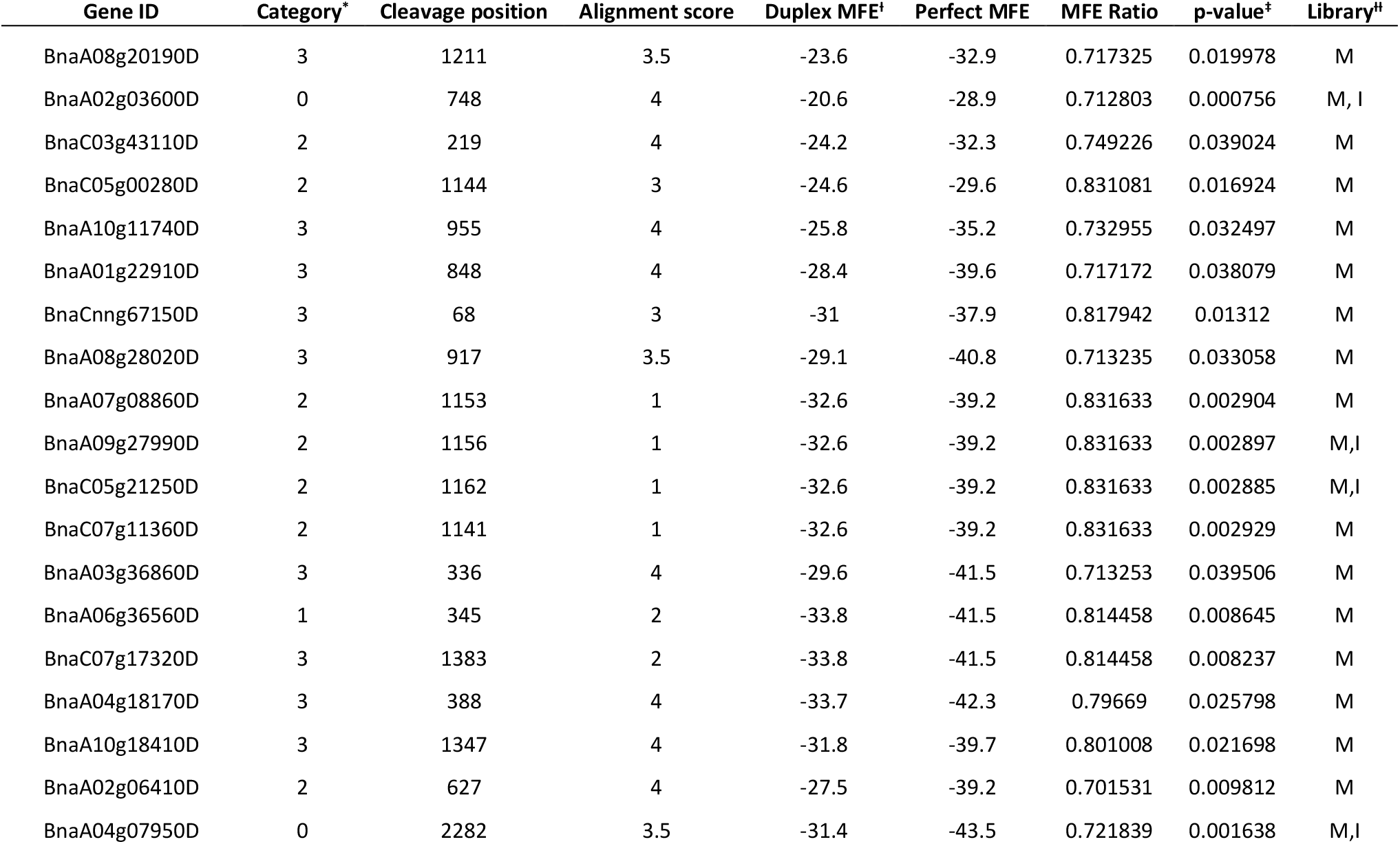
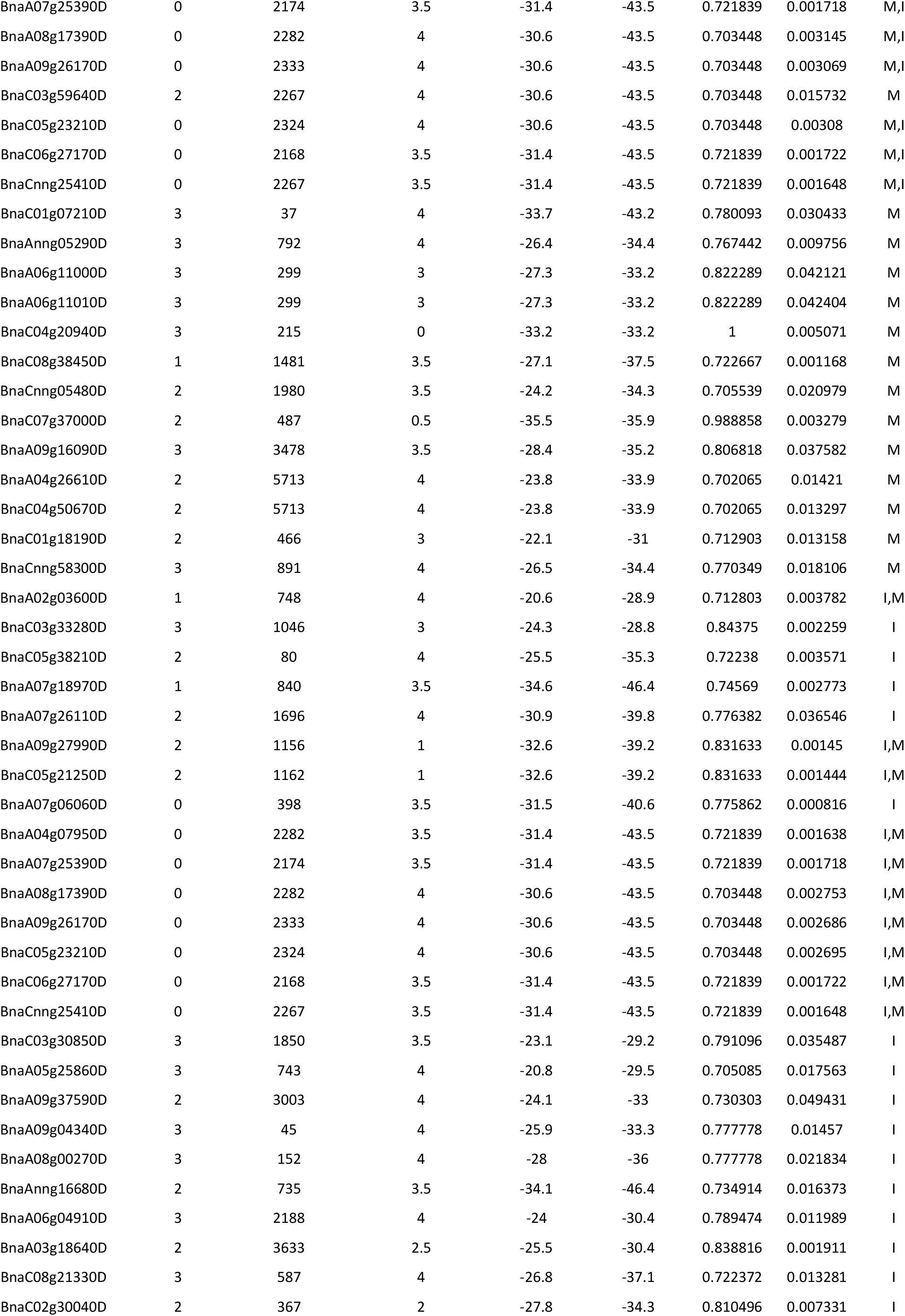
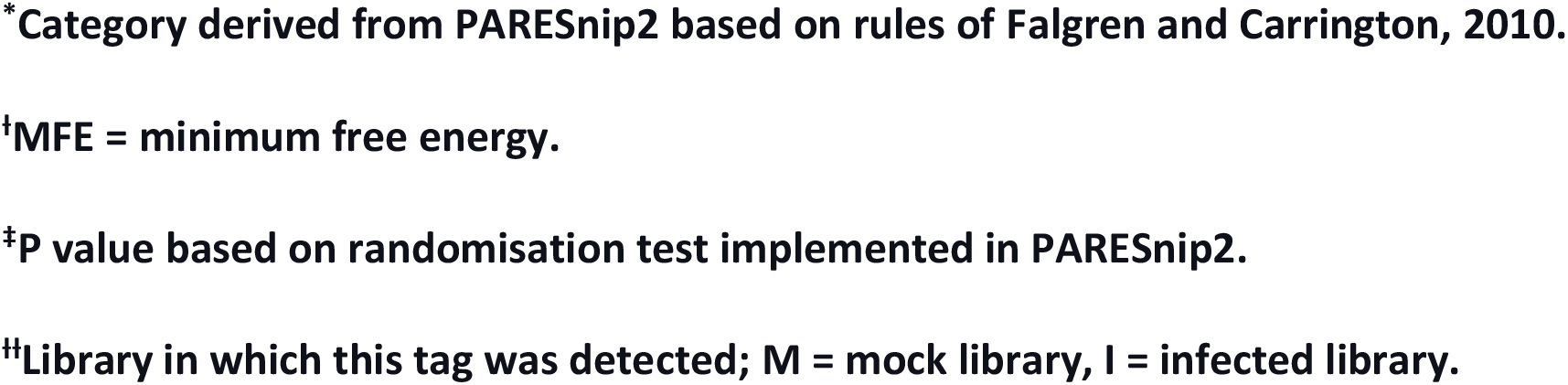
Predicted degraded *Brassica napus* targets of upregulated *B. napus* small RNAs based on degradome sequencing data analysed using PARESnip2.

| Gene ID | Category* | Cleavage position | Alignment score | Duplex MFE<sup>†</sup> | Perfect MFE | MFE Ratio | p-value<sup>‡</sup> | Library<sup>§</sup> |
| --- | --- | --- | --- | --- | --- | --- | --- | --- |
| BnaA08g20190D | 3 | 1211 | 3.5 | -23.6 | -32.9 | 0.717325 | 0.019978 | M |
| BnaA02g03600D | 0 | 748 | 4 | -20.6 | -28.9 | 0.712803 | 0.000756 | M, I |
| BnaC03g43110D | 2 | 219 | 4 | -24.2 | -32.3 | 0.749226 | 0.039024 | M |
| BnaC05g00280D | 2 | 1144 | 3 | -24.6 | -29.6 | 0.831081 | 0.016924 | M |
| BnaA10g11740D | 3 | 955 | 4 | -25.8 | -35.2 | 0.732955 | 0.032497 | M |
| BnaA01g22910D | 3 | 848 | 4 | -28.4 | -39.6 | 0.717172 | 0.038079 | M |
| BnaCnng67150D | 3 | 68 | 3 | -31 | -37.9 | 0.817942 | 0.01312 | M |
| BnaA08g28020D | 3 | 917 | 3.5 | -29.1 | -40.8 | 0.713235 | 0.033058 | M |
| BnaA07g08860D | 2 | 1153 | 1 | -32.6 | -39.2 | 0.831633 | 0.002904 | M |
| BnaA09g27990D | 2 | 1156 | 1 | -32.6 | -39.2 | 0.831633 | 0.002897 | M, I |
| BnaC05g21250D | 2 | 1162 | 1 | -32.6 | -39.2 | 0.831633 | 0.002885 | M, I |
| BnaC07g11360D | 2 | 1141 | 1 | -32.6 | -39.2 | 0.831633 | 0.002929 | M |
| BnaA03g36860D | 3 | 336 | 4 | -29.6 | -41.5 | 0.713253 | 0.039506 | M |
| BnaA06g36560D | 1 | 345 | 2 | -33.8 | -41.5 | 0.814458 | 0.008645 | M |
| BnaC07g17320D | 3 | 1383 | 2 | -33.8 | -41.5 | 0.814458 | 0.008237 | M |
| BnaA04g18170D | 3 | 388 | 4 | -33.7 | -42.3 | 0.79669 | 0.025798 | M |
| BnaA10g18410D | 3 | 1347 | 4 | -31.8 | -39.7 | 0.801008 | 0.021698 | M |
| BnaA02g06410D | 2 | 627 | 4 | -27.5 | -39.2 | 0.701531 | 0.009812 | M |
| BnaA04g07950D | 0 | 2282 | 3.5 | -31.4 | -43.5 | 0.721839 | 0.001638 | M, I |
| BnaA07g25390D | 0 | 2174 | 3.5 | -31.4 | -43.5 | 0.721839 | 0.001718 | M,I |
| BnaA08g17390D | 0 | 2282 | 4 | -30.6 | -43.5 | 0.703448 | 0.003145 | M,I |
| BnaA09g26170D | 0 | 2333 | 4 | -30.6 | -43.5 | 0.703448 | 0.003069 | M,I |
| BnaC03g59640D | 2 | 2267 | 4 | -30.6 | -43.5 | 0.703448 | 0.015732 | M |
| BnaC05g23210D | 0 | 2324 | 4 | -30.6 | -43.5 | 0.703448 | 0.00308 | M,I |
| BnaC06g27170D | 0 | 2168 | 3.5 | -31.4 | -43.5 | 0.721839 | 0.001722 | M,I |
| BnaCnng25410D | 0 | 2267 | 3.5 | -31.4 | -43.5 | 0.721839 | 0.001648 | M,I |
| BnaC01g07210D | 3 | 37 | 4 | -33.7 | -43.2 | 0.780093 | 0.030433 | M |
| BnaAnng05290D | 3 | 792 | 4 | -26.4 | -34.4 | 0.767442 | 0.009756 | M |
| BnaA06g11000D | 3 | 299 | 3 | -27.3 | -33.2 | 0.822289 | 0.042121 | M |
| BnaA06g11010D | 3 | 299 | 3 | -27.3 | -33.2 | 0.822289 | 0.042404 | M |
| BnaC04g20940D | 3 | 215 | 0 | -33.2 | -33.2 | 1 | 0.005071 | M |
| BnaC08g38450D | 1 | 1481 | 3.5 | -27.1 | -37.5 | 0.722667 | 0.001168 | M |
| BnaCnng05480D | 2 | 1980 | 3.5 | -24.2 | -34.3 | 0.705539 | 0.020979 | M |
| BnaC07g37000D | 2 | 487 | 0.5 | -35.5 | -35.9 | 0.988858 | 0.003279 | M |
| BnaA09g16090D | 3 | 3478 | 3.5 | -28.4 | -35.2 | 0.806818 | 0.037582 | M |
| BnaA04g26610D | 2 | 5713 | 4 | -23.8 | -33.9 | 0.702065 | 0.01421 | M |
| BnaC04g50670D | 2 | 5713 | 4 | -23.8 | -33.9 | 0.702065 | 0.013297 | M |
| BnaC01g18190D | 2 | 466 | 3 | -22.1 | -31 | 0.712903 | 0.013158 | M |
| BnaCnng58300D | 3 | 891 | 4 | -26.5 | -34.4 | 0.770349 | 0.018106 | M |
| BnaA02g03600D | 1 | 748 | 4 | -20.6 | -28.9 | 0.712803 | 0.003782 | I,M |
| BnaC03g33280D | 3 | 1046 | 3 | -24.3 | -28.8 | 0.84375 | 0.002259 | I |
| BnaC05g38210D | 2 | 80 | 4 | -25.5 | -35.3 | 0.72238 | 0.003571 | I |
| BnaA07g18970D | 1 | 840 | 3.5 | -34.6 | -46.4 | 0.74569 | 0.002773 | I |
| BnaA07g26110D | 2 | 1696 | 4 | -30.9 | -39.8 | 0.776382 | 0.036546 | I |
| BnaA09g27990D | 2 | 1156 | 1 | -32.6 | -39.2 | 0.831633 | 0.00145 | I,M |
| BnaC05g21250D | 2 | 1162 | 1 | -32.6 | -39.2 | 0.831633 | 0.001444 | I,M |
| BnaA07g06060D | 0 | 398 | 3.5 | -31.5 | -40.6 | 0.775862 | 0.000816 | I |
| BnaA04g07950D | 0 | 2282 | 3.5 | -31.4 | -43.5 | 0.721839 | 0.001638 | I,M |
| BnaA07g25390D | 0 | 2174 | 3.5 | -31.4 | -43.5 | 0.721839 | 0.001718 | I,M |
| BnaA08g17390D | 0 | 2282 | 4 | -30.6 | -43.5 | 0.703448 | 0.002753 | I,M |
| BnaA09g26170D | 0 | 2333 | 4 | -30.6 | -43.5 | 0.703448 | 0.002686 | I,M |
| BnaC05g23210D | 0 | 2324 | 4 | -30.6 | -43.5 | 0.703448 | 0.002695 | I,M |
| BnaC06g27170D | 0 | 2168 | 3.5 | -31.4 | -43.5 | 0.721839 | 0.001722 | I,M |
| BnaCnng25410D | 0 | 2267 | 3.5 | -31.4 | -43.5 | 0.721839 | 0.001648 | I,M |
| BnaC03g30850D | 3 | 1850 | 3.5 | -23.1 | -29.2 | 0.791096 | 0.035487 | I |
| BnaA05g25860D | 3 | 743 | 4 | -20.8 | -29.5 | 0.705085 | 0.017563 | I |
| BnaA09g37590D | 2 | 3003 | 4 | -24.1 | -33 | 0.730303 | 0.049431 | I |
| BnaA09g04340D | 3 | 45 | 4 | -25.9 | -33.3 | 0.777778 | 0.01457 | I |
| BnaA08g00270D | 3 | 152 | 4 | -28 | -36 | 0.777778 | 0.021834 | I |
| BnaAnng16680D | 2 | 735 | 3.5 | -34.1 | -46.4 | 0.734914 | 0.016373 | I |
| BnaA06g04910D | 3 | 2188 | 4 | -24 | -30.4 | 0.789474 | 0.011989 | I |
| BnaA03g18640D | 2 | 3633 | 2.5 | -25.5 | -30.4 | 0.838816 | 0.001911 | I |
| BnaC08g21330D | 3 | 587 | 4 | -26.8 | -37.1 | 0.722372 | 0.013281 | I |
| BnaC02g30040D | 2 | 367 | 2 | -27.8 | -34.3 | 0.810496 | 0.007331 | I |

\*Category derived from PARESnip2 based on rules of Falgren and Carrington, 2010.
<sup>†</sup>MFE = minimum free energy.
<sup>‡</sup>P value based on randomisation test implemented in PARESnip2.
<sup>#</sup>Library in which this tag was detected; M = mock library, I = infected library.

**Figure 3.**
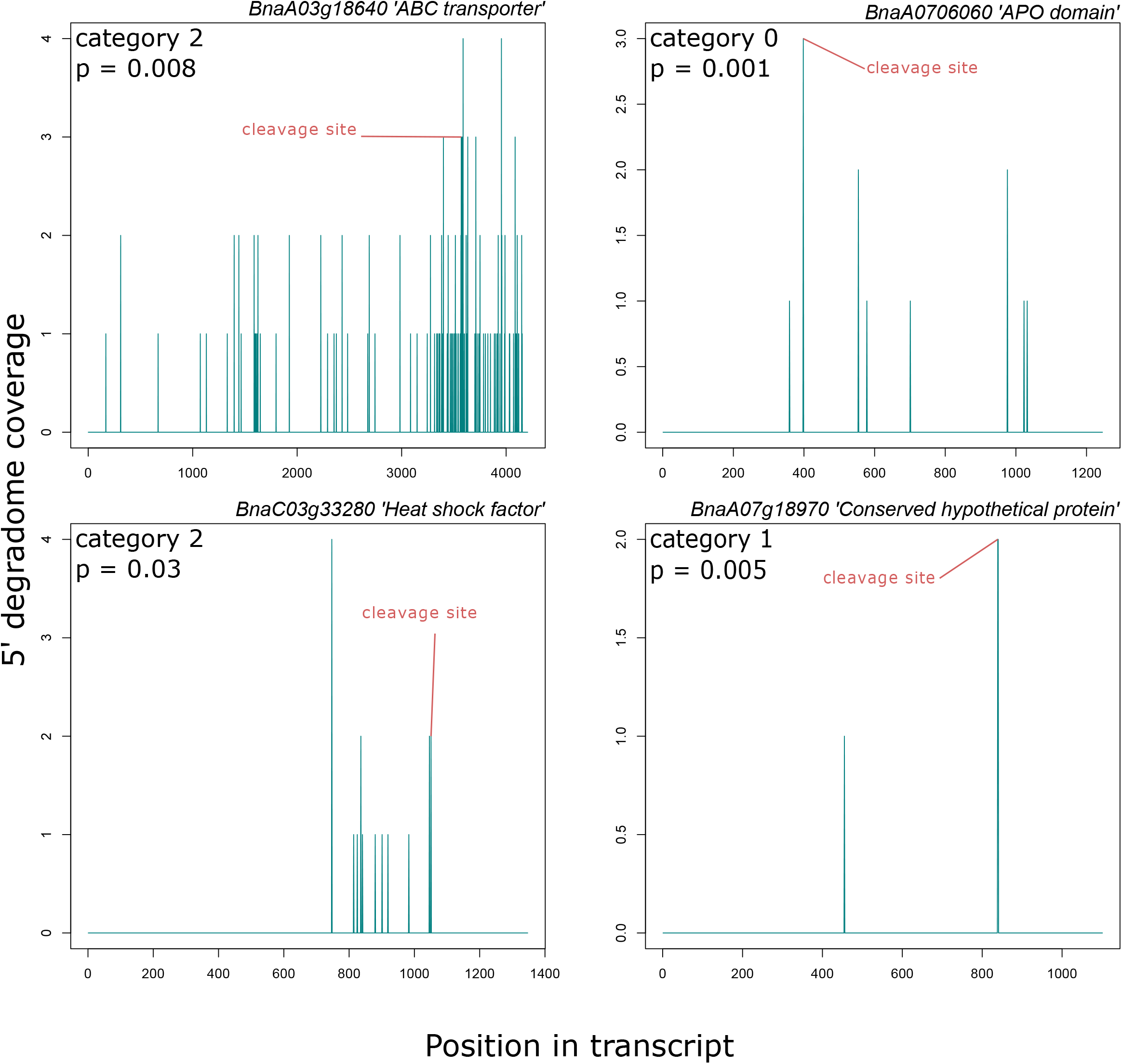
Representative Target plots (T-plots) for infection-specific targets of upregulated sRNAs. The x-axis shows the transcript position in the target genes and the y-axis shows the 5’ read coverage at different positions; cleavage sites predicted by PARESnip2 are labeled in red. The category and *p* value given by PARESnip are shown in the top left hand corners of graphs and the gene IDs and their putative functions are shown above.

### Eight conserved *Brassica napus* miRNAs are differentially expressed in response to *Sclerotinia sclerotiorum* infection

Several miRNAs are evolutionarily conserved in the plant kingdom [53]. We aimed to assess whether conserved *B. napus* miRNAs were expressed in response to *S. sclerotiorum* infection.

Therefore, all six clean libraries were searched against miRBase (Release 22.1). From our libraries, we identified 73 conserved miRNA families with 537 mature miRNA sequences. We found that 61 miRNA families had more than one sequence while 12 miRNA families had only one mature sequence predicted (Fig 4A). Among these miRNA families, miR156 had 42 sequences followed by miR159, and miR166 with 28 and 25 sequences, respectively. Most of these miRNA sequences were 21 nt long, followed by 20, 19 and 18 nt (Fig 4B). There was a 5’ uracil bias in 18-22 nt long miRNA sequences, which agreed with previously reported results in miRNA studies in different plant species (Fig 4C).

**Figure 4.**
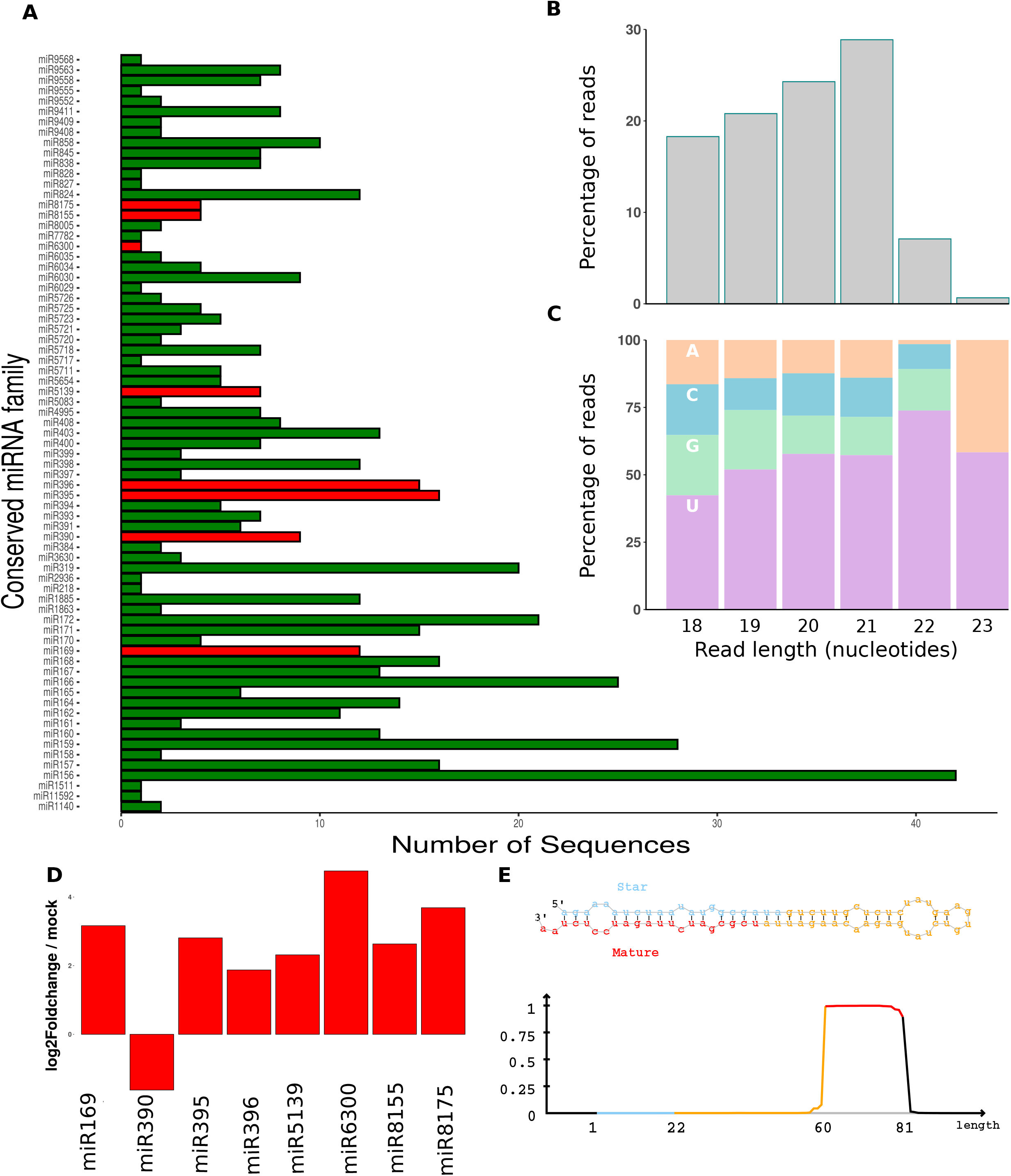
Prediction of infection-responsive microRNAs from the *Brassica napus* genome. **(A)** Histogram of the 73 conserved miRNA families. The y-axis shows the identified conserved miRNA family name and the x-axis shows the number of sequences identified for each miRNA family; red bars are the significantly differentially expressed miRNA families. **(B)** Histogram of read sizes from 537 conserved miRNAs. The y axis depicts the percentage of reads and the x axis read length in nucleotides. **(C)** Histogram of the 537 miRNAs showing percentage of reads (y axis) of each size class (x axis) that had each of the four nucleotides (AGCU) in their 5’ position. **(D)** Eight differentially expressed miRNA families. The x-axis shows the miRNA family and the y-axis shows log2(fold change) of infected samples relative to mock samples. **(E)** The hairpin structure predicted for one of the novel miRNAs predicted by miRDeep2. The red sequence is the mature miRNA and the blue sequence is the passenger strand (miRNA* or ‘star’ sequence); the read coverage for each section of the hairpin is shown in the graph just below the hairpin structure.

Because we had replicated data sets from mock and infected samples, we were able to perform differential expression analysis of the conserved miRNAs. Among the conserved miRNAs, only eight were differentially expressed (Fig 4D). Among them, seven were upregulated after infection and one was downregulated.

Using degradome sequencing, we found that from the 73 conserved miRNA families, 718 and 1,436 cleaved products were obtained from infected and mock libraries, respectively. Four levels of degradome cleavage site confidence, based on read mapping characteristics, are described in [47, 56]. Category 0 is the most confident, followed by categories 1, 2 and 3. Among the 718 targets in infected samples, 507 were in category 0, followed by category 2, 3, and 1 with 90, 87, and 34 targets, respectively based on the abundance of fragment transcripts in the library. Thus, most of the conserved miRNA targets identified with degradome sequencing were of relatively high confidence (Supplementary Table 4) [56]. We found targets for five of the eight differentially expressed miRNA families in the infected sample. Genes containing the term ‘zinc finger domain’ were silenced by miR169, miR390, and miR5139 whereas miR395 and miR396 cleaved transcripts of genes with ‘HCNP-like’ and ‘kinase’ domains.

Several target genes were likely silenced by more than one miRNA family. Among these target genes, 158 and 276 non-redundant transcripts were found in infected and mock samples, respectively. We found 108 transcripts that were common in both degradome libraries; however, 50 targets were unique to only the infected sample so these targets might be infection-induced. Among the 158 targets in infected samples, miR160, miR164, miR167, and miR396 were predicted to target more than 10 genes each. Similarly, miR156, miR6030, miR400, miR393, miR172 and miR171 were predicted to target eight genes (Supplementary Table 5). These miRNAs were reported previously in several studies to regulate gene expression in plants during biotic and abiotic stress [57–60].

### RNA structure-aided prediction algorithms identify 135 novel *Brassica napus* micro RNA loci

After filtering out the exact matches of conserved miRNAs to miRBase, 135 novel miRNA producing loci that did not have any hit in miRBase, were identified from the *B. napus* genome. Among these miRNAs, 67 loci were found to have both passenger strand (miRNA* or ‘star’) and mature strand reads revealing the confidence of these novel miRNAs as per annotation criteria. Fig 4E shows the genomic context of a highly expressed novel miRNA producing locus with the characteristic features attributed to a miRNA origin. A detailed description of the novel miRNA loci is given in Supplementary Table 6.

From our study, we did not find any cleavage signal from novel miRNAs predicted from miRDeep2. We used the same set of miRNAs to predict targets using the psRNA target server. From psRNA target, 12,104 genes were putatively cleaved by these miRNAs (data not shown). Several psRNA targets might be false positives since it was entirely based on a theoretical *in silico* procedure, while a degradome signal is a better reflection of the biological cleavage. It remains to be confirmed whether these novel miRNAs have genuine targets or not.

### Nine *Brassica napus* PHAS loci are differentially expressed in response to *Sclerotinia sclerotiorum* infection

PHAS loci have not been very well characterised in *B. napus*. Therefore, we aimed to identify expressed PHAS loci in the *B. napus* genome from our sequencing data set. We found 26 highly confident PHAS loci in the *B. napus* genome. The genes associated with predicted PHAS loci were annotated by aligning PHAS locus sequences to the NCBI Nucleotide Collection (nr/nt). Among the 26 PHAS genes, about half were related to disease resistance proteins (5 genes), non-coding RNAs (5 genes), and chloroplast related (3 genes). In addition, single genes were found for metal tolerance, pentatricopeptide repeat, cop9 singalosome complex subunit, and photosystem II protein D1. Nine PHAS genes were not homologous to any sequences in NCBI. A total of 182 pha-siRNAs were produced from these loci. Among these siRNAs, 41 were highly expressed, with a read abundance of more than 100 reads.

Since miRNAs are key triggers of pha-siRNA expression, we used the psRNA target server to find the cleavage sites in PHAS loci from the conserved miRNAs we identified. We found six PHAS loci potentially triggered by conserved miRNAs (Table 3). All excised PHAS clusters with their corresponding pha-siRNAs are shown in Supplementary File 1. The miR390-triggered PHAS gene *TAS3* was found to be conserved across different species. In this study, we found two genes possibly targeted by miR390, one of which had sequence similarity to *TAS3* in *A. thaliana*.

**Table 3.**
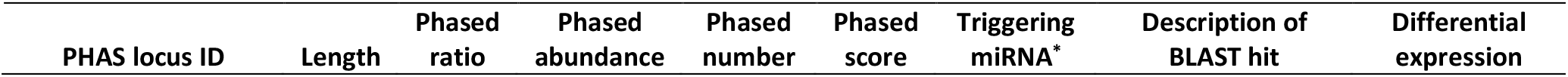
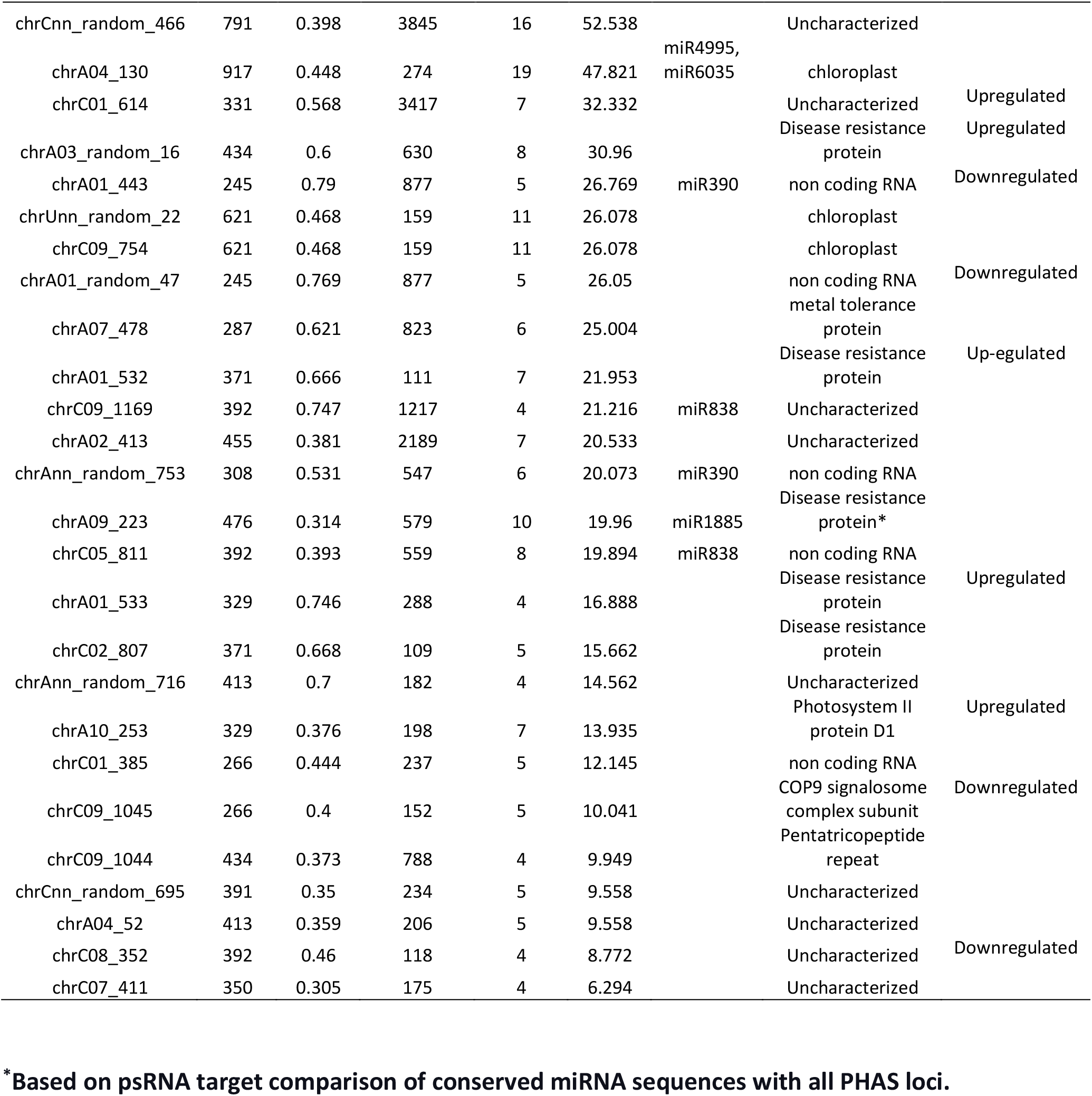
An overview of the characteristics of PHAS loci identified using PHAS tank.

We were able to confirm likely cleavage of one of these loci during infection using degradome sequencing (category 2, P = 0.0019; Fig 5A). We did not find a degradome signal for another five miRNA triggered loci predicted with the psRNA target server. The locus we were able to validate was likely targeted by the conserved miRNA miR1885. Recently, it has been shown experimentally that miR1885 plays a key role in targeting PHAS loci residing within NBS-LRR genes to trigger ta-siRNA production [18]. Accordingly, we found that this locus had homology to NBS-LRR proteins. We identified 10 likely ta-siRNAs produced from this PHAS locus. From the degradome signal, only one of these targets BnaC05g49720D, a galactose oxidase, beta-propeller, had a cleavage signal from degradome sequencing (category 2, P = 0.016; Fig 5B) in the infected sample. Possibly, *B. napus* miRNAs regulate gene expression in response to *S. sclerotiorum* infection through the production of miRNA triggered ta-siRNAs.

**Figure 5.**
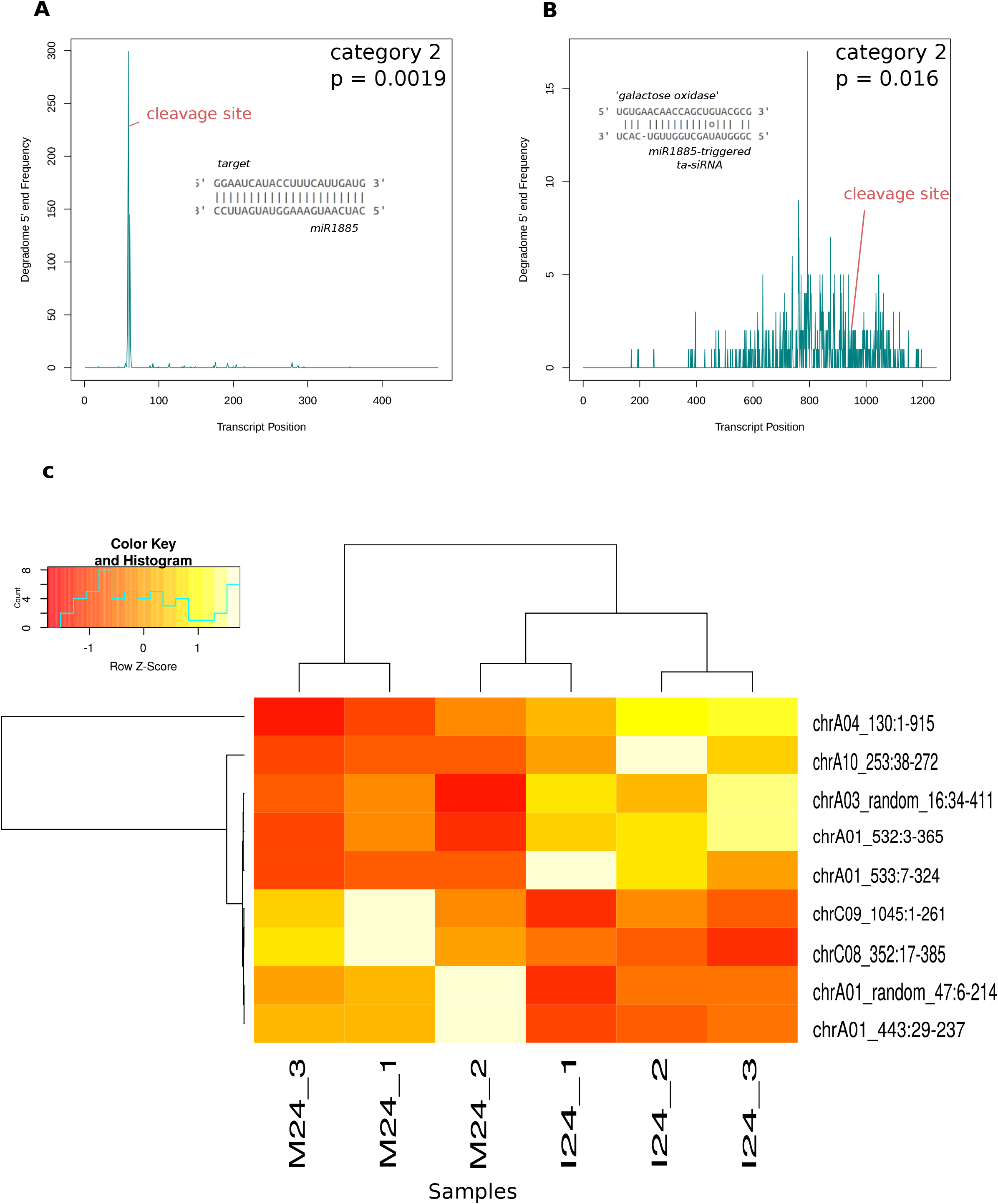
Degradome validation of a TAS gene triggered by miRNA 1885. **(A)** Target plot (T-plot) of the cleavage site on a TAS gene putatively targeted by miRNA 1885. The transcript position in the target gene is on the x-axis and the y-axis shows the 5’ read coverage at each position; the cleavage site predicted by PARESnip2 is labelled in red. The category and *p* value of the PARESnip2 test are given in the top left hand corner of the graph. **(B)** T-plot of cleavage site on a galactose oxidase gene putatively targeted by one of the ta-siRNAs produced by the miRNA 1885-triggered TAS gene. The x-axis shows the transcript position in the target gene and the y-axis shows read 5’ coverage at each position; the cleavage site predicted by PARESnip2 is labelled in red. The category and *p* value from PARESnip2 are given in the top right hand corner of the graph. **(C)** A heat map of 9 differentially expressed PHAS loci plotted with normalized counts from DESEq2.

Differential expression analysis of PHAS loci showed five loci were upregulated and four were downregulated during infection. Among upregulated loci, three were related to disease resistance. The remaining two genes were chloroplast and photosystem II protein D1. Among downregulated loci, two loci were non-coding RNAs, one was related to COP9 signalosome complex subunit and the remaining one was not characterized. Fig 5C shows a heat map of 9 differentially expressed PHAS loci.

To gain a global overview of genes targeted by pha-siRNAs we used the psRNA target server to find the targets of 41 highly expressed pha-siRNAs. We found 5,918 transcripts that might be regulated by this class of sRNA. We did GO term enrichment analysis of these targets and found regulation of several biological processes (Supplementary Table 7). The terms ‘posttranscriptional gene silencing’ (GO:0035194), ‘cellular potassium ion homeostasis’ (GO:0030007), ‘regulation of ARF protein signal transduction’ (GO:0032012) and ‘threonyl-tRNA amino acylation’ (GO:0006435) were significantly enriched while the ‘terms oxidation-reduction process’ (GO:0055114), and ‘regulation of transcription DNA-templated’ (GO:0006355), ‘carbohydrate metabolic process’ (GO:0005975) were depleted.

We also specifically investigated the targets of the miR1885-triggered ta-siRNAs. We found 1,601 targets of these sRNAs with psRNA target. GO term enrichment analysis showed that these ta-siRNAs possibly regulate protein phosphorylation (GO: 0006468), transcription factors (GO:0045944), vesicle mediated transport proteins (GO:0016192), and fucose metabolic pathway genes (GO:0006004) (Supplementary Table 8).

### Further validation of targeting using 5’ rapid amplification of cDNA ends

5’ RACE was used as a validation technique to find putative cleavage sites in the sRNA target gene *BnA01g27570D*, which is an ethylene response factor. This gene was chosen as it is possibly targeted by a novel sRNA from a locus identified by ShortStack. This locus is not conserved or characterised and neither a pha-siRNA nor a miRNA. Fig 6A shows the 5’-RACE product of the predicted cleavage site from the infected sample. The complementary sRNA region in the mRNA and the validation of the cleavage point through degradome reads is shown in Fig 6B. A T-plot of the cleavage signal is shown in Fig 6C.

**Figure 6.**
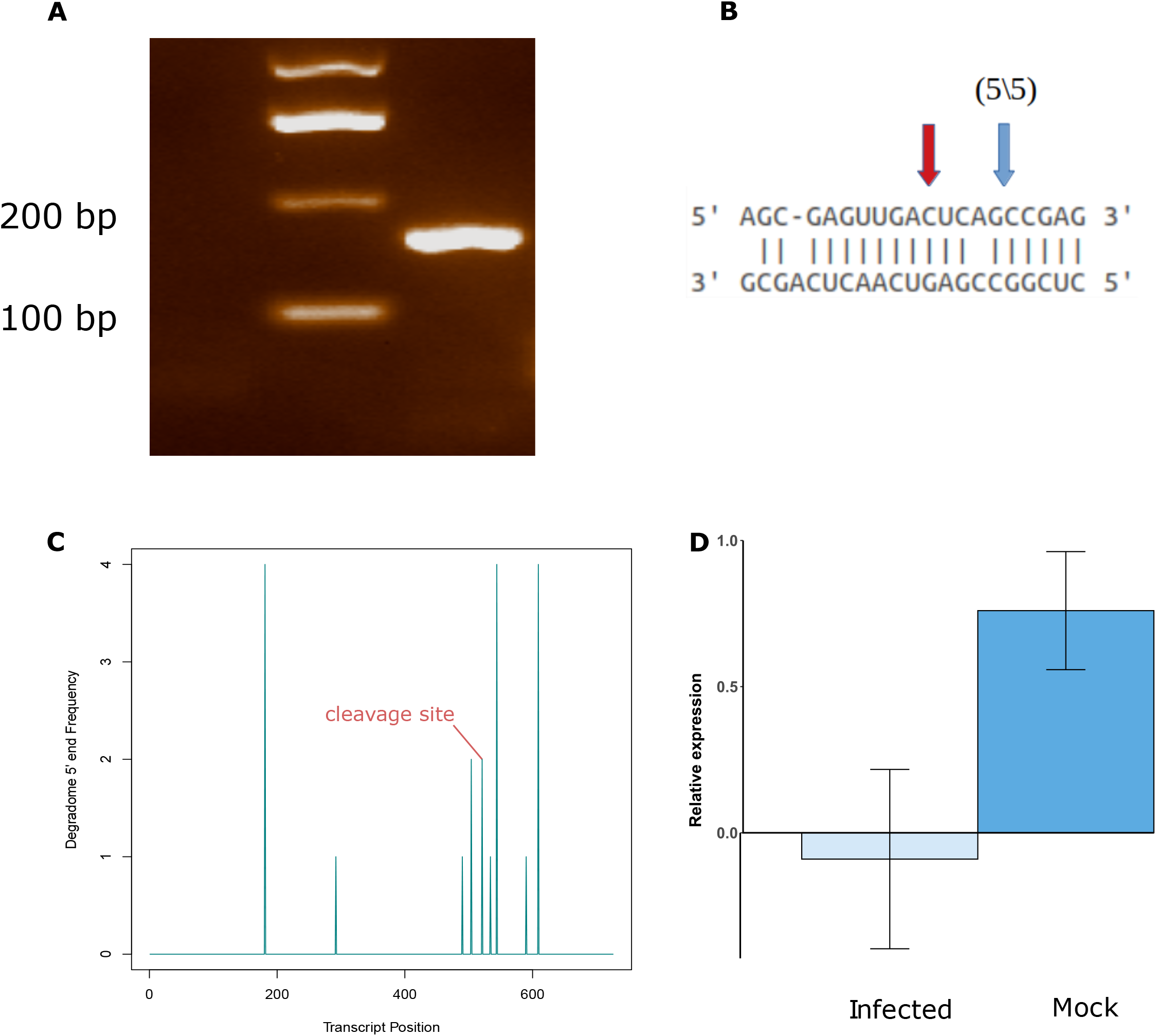
5’ Rapid amplification of cDNA ends (RACE), degradome result, and qPCR for an ethylene response factor gene putatively cleaved by a novel sRNA. **(A)** Gel electrophoresis of the 5’-RACE result showing a band of the correct size. **(B)** Sequence complementarity of the sRNA and its target. The blue arrow shows the cleavage site identified from sequencing the 5’-RACE product and the red arrow shows the cleavage site identified with degradome sequencing. **(C)** Target-plot (T-plot) of the degradome result of the 5’-RACE validated gene showing the transcript position in the target gene (x-axis) and read 5’ coverage at each position (y-axis); the cleavage site predicted by PARESnip2 is labelled in red. **(D)** qPCR result of target gene in mock and infected sample (x-axis) and relative expression of gene to the house keeping actin gene.

We also assessed the expression of this target gene during infection using qPCR. The average log (2^−ΔCt^) value calculated for mock and infected sample was 0.76 and −0.09 respectively with a standard deviation of 0.35 and 0.53 Fig 6C. The combined degradome, 5’RACE, and qPCR results showed that a novel *B. napus* sRNA likely regulates an ethylene response factor gene during *S. sclerotiorum* infection.

## Discussion

Small RNA-omics studies have revealed the tight regulation of host immune pathways in various studies [19, 55, 61]. From our study, we identified different classes of sRNAs in *B. napus* plants that responded to infection with *S. sclerotiorum* and showed how they may be involved in regulating different sets of genes using degradome sequencing.

### Biogenesis of small RNA in *Brassica napus* is altered upon *Sclerotinia sclerotiorum* infection

The distribution of size classes of sRNAs was different between mock and infected samples. Size class of sRNAs gives insights into their biogenesis, for example 21 nt sRNAs are processed by DCL1 and DCL4, whereas 22 nt are formed through the action of DCL2; 24 and 26 nt sRNAs are formed by DCL3 [62]. Similarly, sRNAs with 5’ uracil, adenine or cytosine bias are loaded into AGO1, AGO2 and AGO4 or AGO5, respectively [63]. Furthermore, it has been reported previously that variation in sRNA lengths also has effects on downstream function of sRNAs [64]. Overall, our data suggest that a different set of sRNA biogenesis pathways are initiated upon infection with *S. sclerotiorum*.

### Differential expression analysis showed that a large number of *B. napus* small RNA generating loci were upregulated in response to challenge with *S. sclerotiorum*

A total of 730 sRNA loci were upregulated in *B. napus* in response to *S. sclerotiorum* infection. The upregulated sRNAs mostly belonged to the size classes of 20 and 21 nt with a 5’ bias of uracil, whereas downregulated sRNAs were overrepresented for 24 nt sequences with varied 5’ biases expression pattern than upregulated sRNAs. This suggests that upon pathogen infection, the host recruits different Dicer and Argonaute proteins for subsequent gene silencing. The upregulated sRNAs silenced many genes related to stress signaling as determined from degradome tags. This suggests that the enhancement of disease symptoms might be accompanied with the negative regulation of plant immune genes during *S. sclerotiorum* infection of *B. napus*.

### Identification of conserved miRNAs and their regulatory cascade

The miRNAs are sRNAs which are produced from short hairpin precursors. Based on these criteria, thousands of miRNAs have been identified and deposited in miRBase from many plants. The majority of these miRNAs show high conservation. However, *Brassica*-specific miRNAs discovered so far are less numerous than those of *A. thaliana*, *M. truncatula*, and *O. sativa* revealing several miRNAs are yet to be discovered in this species.

Jian et al. (2018) identified 30 conserved miRNA families with 77 known miRNAs during *S. sclerotiorum* infection of *B. napus* [31]. In this study we identified more than double this number of conserved miRNA families. This might be due to the sequencing depth in our study and our use of replicated samples. Moreover, 135 novel miRNA loci were discovered with 67 highly confident loci that had both mature and star (passenger strand) read counts.

Some of the conserved miRNAs were upregulated in response to pathogen attack and were previously reported to have a role in regulating plant immunity. Expression of a large number of transcription factors and auxin signaling pathway genes was likely regulated by these miRNAs. Some of these miRNAs had multiple targets in specific classes. For example, miR167 targets auxin response factors, miR62 has target sites in Argonaute and Dicer proteins, miR396 silences protein kinases, and miR6030 and miR1885 have target sites on disease resistance proteins. These findings suggest that miRNAs are involved in regulation of multiple aspects of the immune response of *B. napus* to *S. sclerotiorum*.

Previous studies showed that decreased expression levels of miR396 over the infection period rendered hosts more resistant to fungi [55]. In our study, miR396 was upregulated in the infected samples suggesting that it could possibly enhance disease symptoms. Degradome data suggested that this miRNA regulates a protein kinase, which could be a key part of immune signaling in *B. napus*. Rice miRNA 169 negatively regulates rice immunity against *M. oryzae* via silencing of nuclear factor Y-A. In our study, miR169 was upregulated in the infected samples and possibly silenced a zinc finger, Sec23/Sec24-type transcription factor [61]. Overexpression of miR400 in *A. thaliana* made the plant more susceptible to necrotrophic fungi [65]. Therefore, overexpression of some miRNAs can enhance the disease symptoms by suppressing plant immunity-related genes.

### Identification of 26 PHAS genes from this study with six loci putatively triggered by conserved miRNAs

With the aid of the degradome library we showed that miR1885 can trigger a disease resistance *TAS* gene which subsequently produces 10 ta-siRNAs for gene regulation. Identification and elucidation of the regulatory network of pha-siRNAs is important so that the expression of these phasiRNAs can be controlled by changing the expression of their miRNA triggers. This strategy could be useful to modulate the degree of silencing of endogenous and exogenous target genes.

An integrated analysis revealed that a *B. napus* sRNA regulates an ethylene response factor gene during pathogen attack. Ethylene response factors regulate several jasmonate (JA) and (ET) pathways and are key players in plant innate immunity [50]. Our combined degradome, 5’ RACE and qPCR results showed that the expression of one of the ethylene response genes is suppressed in *B. napus* after *S. sclerotiorum* infection. The silencing of this gene was mediated by a novel sRNA which was not characterized before. The ARF protein signal transduction was previously reported to be regulated by miR390 triggered tasiRNAs [9].

## Conclusions

In conclusion, our comprehensive data set allowed us to investigate overall pathogen-responsive RNA interference-based regulation of host transcripts and the actions of specific small RNA classes in response to pathogen attack. Our data suggest that targets of general RNA interference may be associated with auxin and jasmonic acid signaling pathways. *B. napus* plants may differentially express both pha-siRNAs and conserved miRNAs when challenged with a necrotrophic pathogen.

## Methods

### Biological materials

An Australian *S.sclerotiorum* isolate (CU8.24) originally collected from South Stirling WA was used for infection assays [32]. Mature sclerotia were cut into halves and placed onto 9 cm Petri dishes containing potato dextrose agar (PDA). After germination from the sclerotium, mycelium was subcultured onto fresh PDA medium. After 48 hours of incubation, mycelial plugs were placed on fully expanded second or third leaves of one-month-old *B. napus* plants (AV Garnet). The plants were grown for a month in a growth chamber with 16 h of daylight and 8 h of darkness. After infection, plants were carefully covered with a polythene bag to increase the humidity, thereby facilitating the infection process. Twenty-four HPI, a characteristic necrotic lesion was observed on the infected leaves. The infected tissues were carefully excised using sterilized scissors and immediately frozen in liquid nitrogen and stored at −80 °C until RNA extraction for sequencing. For mock samples, PDA only agar plugs were used without any fungal mycelium. Three leaves from three different plants were pooled together for each replicate. For small RNA sequencing, three biological replicates were sequenced separately while 2 degradome libraries were sequenced by pooling all infected replicates as one library and all mock replicates as another library.

### Total RNA extraction and sequencing

Total RNAs were extracted using the TRIZOL reagent following the manufacturer’s protocol (Invitrogen Carlsbad, CA, USA). After extraction, total RNAs were quantified using a Nanodrop spectrophotometer, and Qubit. The integrity of RNA samples was checked using agarose gel electrophoresis. Three to five μg and 25-30 μg of total RNA were sent to Novogene (Singapore) for small RNA and degradome sequencing respectively. The sRNA sequencing was done using the NEBNext^®^ Multiplex Small RNA Library Prep Kit for Illumina^®^ with single end 50 bp reads according to the manufacturer’s protocol. Degradome sequencing was done as mentioned in [33]. In brief, the construction of a degradome library was started from the degradation site (with monophosphate group) of the degraded mRNA. The sequencing adaptors were added to both ends of the degradation library and a library size of around 200-400 bp was selected. The sequencing was performed on a Hiseq 2500 SE50.

### Analysis of small RNA sequencing data

Raw reads were trimmed using cutadapt software (version 1.15) optimized for single-end reads with a setting of cutadapt-a (universal True Seq adapter) −m18 −M30 [34]. The quality of filtered reads was checked by running in FastQC [35]. Reads with a length in the range of 18-30 nt were retained. Trimmed infected reads were assigned to the fungal [36] and plant [37] reference sequences using bbsplit software in the bbmap [38] program keeping ambig2 option set as toss. The option ambig2 removes all the reads that map to both references with equal confidence. The reads that were unique to the *B. napus* genome were kept for prediction of *B. napus* sRNAs.

For prediction of *B. napus* sRNA biogenesis loci, clean reads were aligned to the reference genome of *B. napus*. We used ShortStack [39] to gain an overall idea of sRNA producing loci from the *B. napus* genome and to characterize highly expressed sRNA loci after infection. Each library was used as a single entity without collapsing for input into ShortStack. For conserved miRNA prediction, we matched the clean reads against miRBase (version 22) (http://www.mirbase.org) using the miRProf program in UEA small RNA workbench [40]. The reads that matched to mature miRNAs in the miRbase database with 0 mismatches were considered as conserved miRNAs. The remaining reads that did not match miRBase were parsed for the prediction of novel miRNA-producing loci using the miRDeep2 program [41]. Differential expression analysis was done using the Bioconductor package DESeq2 in R with estimate variance – mean dependencies. We used the raw read counts of 6 individual libraries from ShortStack to find differentially expressed sRNAs. The sRNAs with a Benjamini-Hochberg corrected p-value of < 0.05 were considered as differentially expressed sRNAs [42].

The phasing patterns of loci were predicted with a Perl script from the PHAS tank software (version 1.0) [43]. To find the miRNA triggered phased initiator loci, complementary cleavage sites of predicted miRNAs on PHAS loci were searched using the psRNA target server assuming that the 10^th^ nucleotide position on the miRNA is a cleavage start position of its targeted PHAS loci [14].

### Analysis of sRNA targeting using *in silico* target prediction and degradome sequencing data

To determine whether reads originated either from the plant or the fungus we used bbsplit to categorise infected degradome reads as fungal or plant-specific reads. The filtered reads were separated from potential structural RNAs by filtering against the RFAM database [44] using the program Infernal (version 1.1.3) [45].

To gain an overall view of gene regulation by plant sRNAs in mock and infected samples, we used all valid non-redundant sRNAs to find their degradome tags in mock and infected samples with the program PARESnip2 [47]. To gain a more detailed understanding of silencing by different sRNA classes, we used four different datasets: the one highly expressed major RNA per locus from the ShortStack program, the conserved miRNAs from miRbase, novel miRNAs annotated from miRDeep2, and ta-siRNAs produced by predicted PHAS loci.

We used either the psRNA target server with a default setting and an expectation score of 5 for computational prediction of sRNA targets [46] or PARESnip2 [47] to validate the cleavage sites from Degradome datasets following the rules of Fahlgren and Carrington [48]. We retained the targets with category number 0-3 as mentioned in [47]. Category-0 are targets with a degraded products having degradome peaks more than one read and the maximum on the transcript where there is only one maximum. Category-1 are those having degradome peaks greater than one read and are the maximum on the transcript, but there is more than one maximum. Category-2 peaks are those that have reads more than one and are above the average fragment abundance on the transcript. Category-3 signals are those that have greater than one read and are below or equal to the average fragment abundance on the transcript. Further verification of PHAS locus activation by specific miRNAs was also investigated using the degradome sequencing tags with PARESnip2.

### Gene ontology enrichment analysis

Gene ontology enrichment analysis was conducted on sRNA target transcripts with the topGO program from R 3.6.1 Bioconductor package. GO term enrichment tests were performed separately on mock and infected samples. In each case, the background set was all GO terms in the *B. napus* genome and the foreground set was any gene with evidence of degradome targeting. The mock and infected samples were compared to identify genes that were enriched in the mock and depleted in the infected sample or vice versa. GO terms with a p-value < 0.05 were considered as significantly enriched or depleted [49].

### Five prime rapid amplification of cDNA ends of a cleaved target

We conducted a 5’-RACE experiment on one of the ethylene response factor genes from our degradome dataset that is potentially cleaved by a plant sRNA in the infected sample but not in the mock sample. The reason for choosing this gene is directed by previous pieces of literature where these classes of genes were shown to be crucial for defence responses in plants against pathogen attack [50]. Furthermore, the sRNA targeting this gene was hitherto uncharacterised, and it is not a miRNA or phasiRNA. We used two independent samples collected from independent infection assays to conduct 5’ RACE using the first choice RACE kit following the manufacturer’s protocol (Applied Biosystems, USA) without adding calf intestinal Phosphatase enzyme. One sample was the same as the one used for degradome sequencing while the other was not.

In brief, 5’ RACE adapters were ligated to 5 μg of total RNA, which was reverse transcribed using the universal RT primer provided in the kit and the MMLV transcriptase. The first PCR was conducted on 1 μL of cDNA with a 5’ outer RACE primer and gene-specific outer primer. The second nested PCR was done using the first PCR product with a 5’ inner RACE primer and inner nested PCR primers. The PCR product was visualized on a 2% Agarose gel. The amplified DNA fragment was gel purified and cloned into TOP TA vector and 5 independent clones were Sanger sequenced.

### Quantitative polymerase chain reaction

The expression levels of the RACE validated target gene were analysed by qPCR. One to five ug of total RNAs from mock and infected *B. napus* leaf samples were converted to cDNA using the MMLV reverse transcriptase kit (Sigma-Aldrich). The cDNA samples were then diluted 1/20 before qPCR. The qPCR analysis was performed using the Bio-Rad Taq Universal SYBR Green Supermix according to the manufacturer’s instructions. The thermocycler settings were 95°C for 2min, then 95°C for 15sec, 60°C for 30sec and 72°C for 15sec, and cycled for 40 times, followed by 72°C for 2min. Three biological and three technical replicates were used for each sample. Relative expression was calculated as per log(2^−ΔCt^) method normalized to the *B. napus* housekeeping actin gene. The primers and adapters used for 5’ RACE and qPCR experiments are listed in Supplementary Table 1.

## Declarations

### Ethics approval and consent to participate

Not applicable.

### Consent for publication

Not applicable.

### Availability of data and materials

The smallRNA and degradome sequencing data has been deposited in GenBank under BioProject PRJNA678586

### Competing interests

The authors declare no conflict of interest.

### Funding

This work was undertaken within the Centre for Crop and Disease Management (CCDM), a co-investment between Curtin University and the Grains Research and Development Commission (GRDC project number CUR00023). RR was funded by scholarships from the Australian Government Research Training Program and the Commonwealth Scientific and Industrial Organisation (CSIRO).

### Authors’ contributions

RR designed and analysed experiments and wrote the first manuscript draft. TEN and LGK were involved in high level and technical experimental discussions and helped develop the manuscript. MCD helped design, analyse and present experimental data, oversaw experimentation and co-wrote the manuscript. All authors read and approved the final version of the manuscript.

## Acknowledgements

This work was supported by resources provided by the Pawsey Supercomputing Centre with funding from the Australian Government and the Government of Western Australia. Authors would like to thank Dr. Yuphin Khentry for providing technical assistance.

## Notes

### Competing Interest Statement

The authors have declared no competing interest.

